# A Dimer between Monomers and Hexamers – Oligomeric Variations in Glucosamine 6-Phosphate Deaminase Family

**DOI:** 10.1101/2022.07.06.499027

**Authors:** Sathya Srinivasachari, Vikas R. Tiwari, Tripti R. Kharbanda, Ramanathan Sowdamini, Ramaswamy Subramanian

## Abstract

In bacteria that live in hosts whose terminal sugar is a sialic acid, Glucosamine 6-phosphate deaminase (NagB) catalyzes the last step in the conversion of sialic acid into Fructose-6-phosphate, which enters the glycolytic pathway. The enzyme exists as a hexamer in Gram-negative bacteria and is shown to be allosterically regulated. In Gram-positive bacteria, it exists as a monomer and lacks allosteric regulation. Our identification of a dimeric Gram-negative bacterial NagB motivated us to characterize the structural basis of the various oligomeric forms. We characterized the crystal structures of NagB from two Gram-negative pathogens, *Haemophilus influenzae (Hi)* and *Pasturella multocida (Pm)*. The *Hi*-NagB is active as a hexamer, while *Pm*-NagB is active as a dimer. We confirm that this is not a crystallographic artifact by cryo-electron microscopy. Both Hi-NagB and Pm-NagB contain the C-terminal helix, and the residues in the interface involved in oligomerization are conserved. The hexamer is described as a dimer of trimers. In the Pm-NagB dimer, the dimeric interface is conserved. This would suggest that the three possible oligomeric forms of NagB are a monomer, a dimer, and a trimer of dimers. Computational modeling and MD simulations indicate that the residues at the trimeric interface have less stabilizing energy of oligomer formation than those in the dimer interface. We propose that Pm-NagB is the evolutionary link between the monomer and the hexamer forms.

## 1 INTRODUCTION

Glucosamine 6-phosphate deaminase (NagB) is an aldose-ketose isomerase that is the final step of the sialic acid catabolic pathway in the tightly regulated and conserved *nag-nan* operon. It converts D-glucosamine 6-phosphate to D-fructose-6-phosphate (F-6-P) and ammonia. In its hexameric form, the enzyme is allosterically activated by N-acetylglucosamine-6-phosphate (GlcNAc6P)[1]. The monomeric form found in Gram-positive bacteria does not show allostery[2,3]. F-6-P then enters the glycolytic cycle as a carbon source for the bacteria to survive. Sialic acids are nine carbon acidic sugars universally present on all mammalian cell surfaces as terminal sugars of glycoproteins and glycolipids[4]. Bacteria colonizing the heavily sialylated mammalian gut and respiratory tract evolved a unique mechanism for scavenging the sialic acid from the host and using it as an energy source for survival[4]. Apart from this, few opportunistic pathogens have found ways to sugar-coat their cell surfaces with sialic acid and use it for molecular mimicry, thereby evading human defense mechanisms[5][6,7]. Several reports show that the complex regulatory interplay of the sialometabolic genes significantly influences the colonization and pathogenicity of commensal pathogens[5]. Sialyation is known as a major virulence determinant in the pathogenesis of biofilm formation *in H. influenzae* and *P. multocida. H. influenzae* causes otitis media, and *P. multocida* is zoonotic and causes various diseases in animals and birds[4,6].

Studies in *S. mutans* show that NagB inactivation leads to a decrease in the expression of virulence factors and also impedes biofilm formation and saliva-induced aggregation [7]. Similarly, NagB deletion mutants in *S. pneumoniae* and *B. subtilis* indicate that NagB is important for growth using sialic acid as the sole carbon source[8]. These suggest the critical role that NagB plays in pathogen survival. Structures of NagB from Gram-positive bacteria, *E.coli*, and human enzymes have been reported [9,10]. The structures fall into two classes hexameric (dimer of trimers as described by earlier authors), enzymes that are allosterically regulated, and monomeric enzymes. This paper focuses on investigating the structure of the deaminases from *H. influenza (Hi)* and *P. multocida (Pm)* (both Gram-negative bacteria), where Hi NagB is a hexamer. Despite very high sequence similarity, conserved active site, and the presence of the C-terminal helix, Pm NagB is a dimer (Figure 1) [10,11].

**FIGURE 1.**
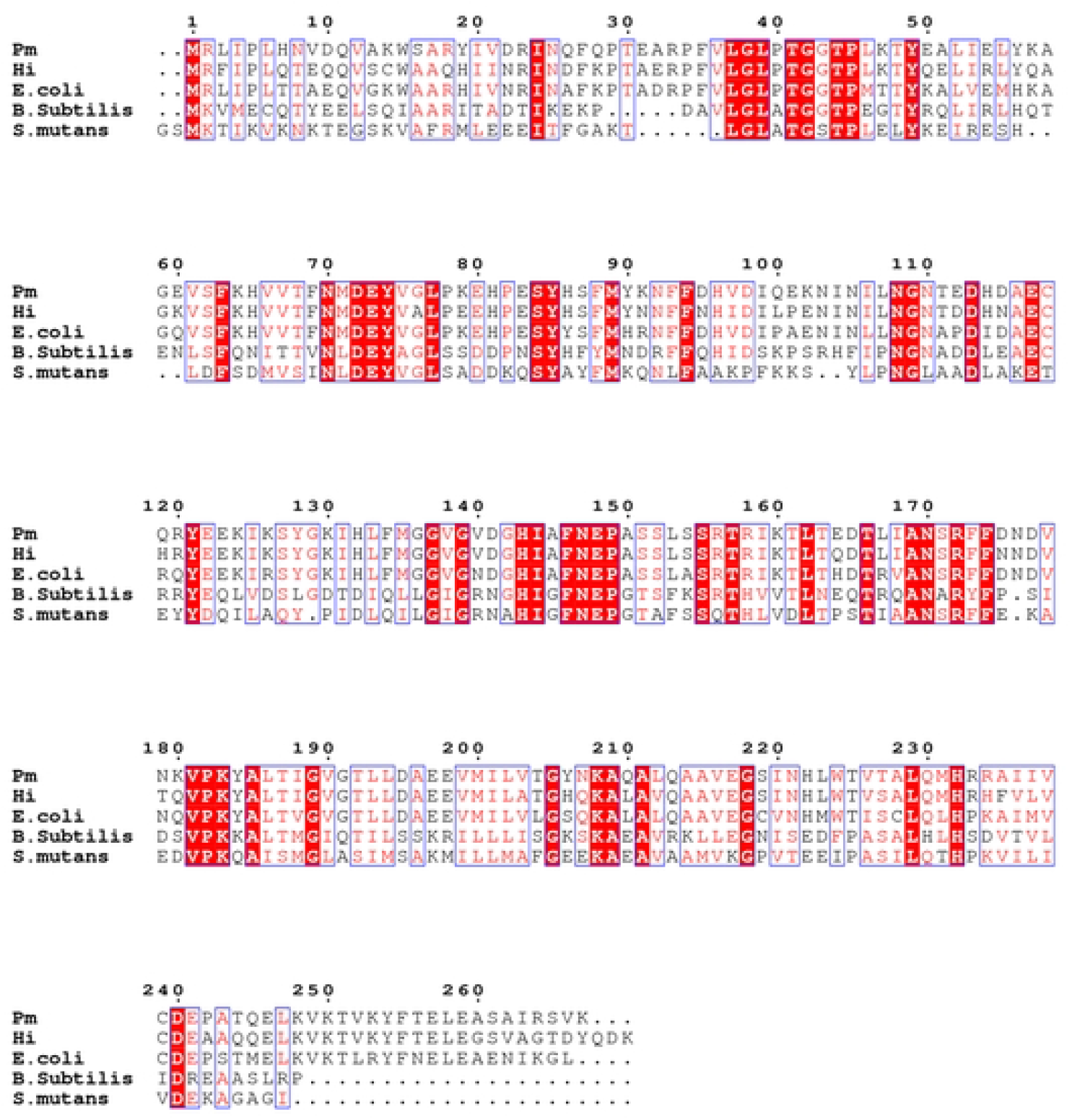
Multiple sequence alignment of NagB from *E.coli, Pm* and *Hi, B. subtilis* and *S.mutans* generated using ESPript 3.0. The secondary structural elements are marked based on the structure of Pm-NagB.

## 2 MATERIALS AND METHODS

### 2.1 Protein Expression and Purification (*Hi*NagB and *Pm*NagB)

Proteins were expressed from the recombinant plasmids, synthesized using gateway cloning technology[12], in *Escherichia coli* BL21(DE3)* cells. The cells were grown in an LB medium containing ampicillin at 37 °C to an OD_600_ 0.6. The cells were then induced with 0.3 mM IPTG. After induction, the cells were allowed to grow for three hours at 37 °C. The cells were harvested and centrifuged at 6000 rev min ^-1^ for 30 mins at 4 °C. Each 1 L cell pellet was resuspended in 25 mL of resuspension buffer (20 mM HEPES, 150 mM NaCl, 5 mM Imidazole, pH 8.0) with a protease inhibitor cocktail without EDTA (Roche). The cells were lysed using an emulsiflex at 100 MPa. The lysate was centrifuged at 1300 rev min^-1^ for 45 mins. The supernatant was purified using Ni-NTA affinity chromatography. His-tagged *Hi*NagB and *Pm*NagB were eluted using a buffer solution containing 20 mM HEPES buffer, pH 8.0, 150 mM NaCl, 300 mM Imidazole. The proteins were further purified using size exclusion chromatography using S200 superdex column with a buffer solution of 20 mM Sodium phosphate, pH: 7.4, 50 mM Sodium chloride, and 10 % glycerol. Purified proteins were concentrated to 15 – 16 mg/mL for crystallization.

### 2.2 Crystallization, Data collection, and Processing

Hanging-drop vapor-diffusion experiments were performed using a Mosquito robot (TTP Labtech). Crystals of *Pm*NagB and *Hi*NagB were obtained by mixing 0.5 ml screening solution with 0.5 ml of protein solution (15-16 mg/mL) and equilibrating the mixture against 100 ml of commercially available crystallization screen conditions (Crystal Screen, Hampton Research). Rod-shaped crystals appeared for PmNagB within 15 days at 18° C in the presence of 100 mM Sodium Cacodylate/HCl pH 6.5, 200 mM Magnesium Chloride, and 20 % PEG 1000. Cuboid-shaped crystals appeared for HiNagB within 15 days at 18° C in the presence of 2 % w/v Tasciminate pH 5.0, 0.1 M Sodium Citrate tribasic pH 5.6, 16 % PEG 3350 with 0.1 M Strontium Chloride. 6H_2_O as additive.

Diffraction data were collected to 2.3 Å resolution from a single crystal of *Pm*NagB and 3.0 Å resolution from a single crystal of *Hi*NagB on the PROXIMA-1 beamline at the SOLEIL synchrotron source. Data were processed with XDS/XSCALE[13] and scaled with AIMLESS from the CCP4 suite [14]. The data-processing statistics are provided in Table 1.

**TABLE 1:**
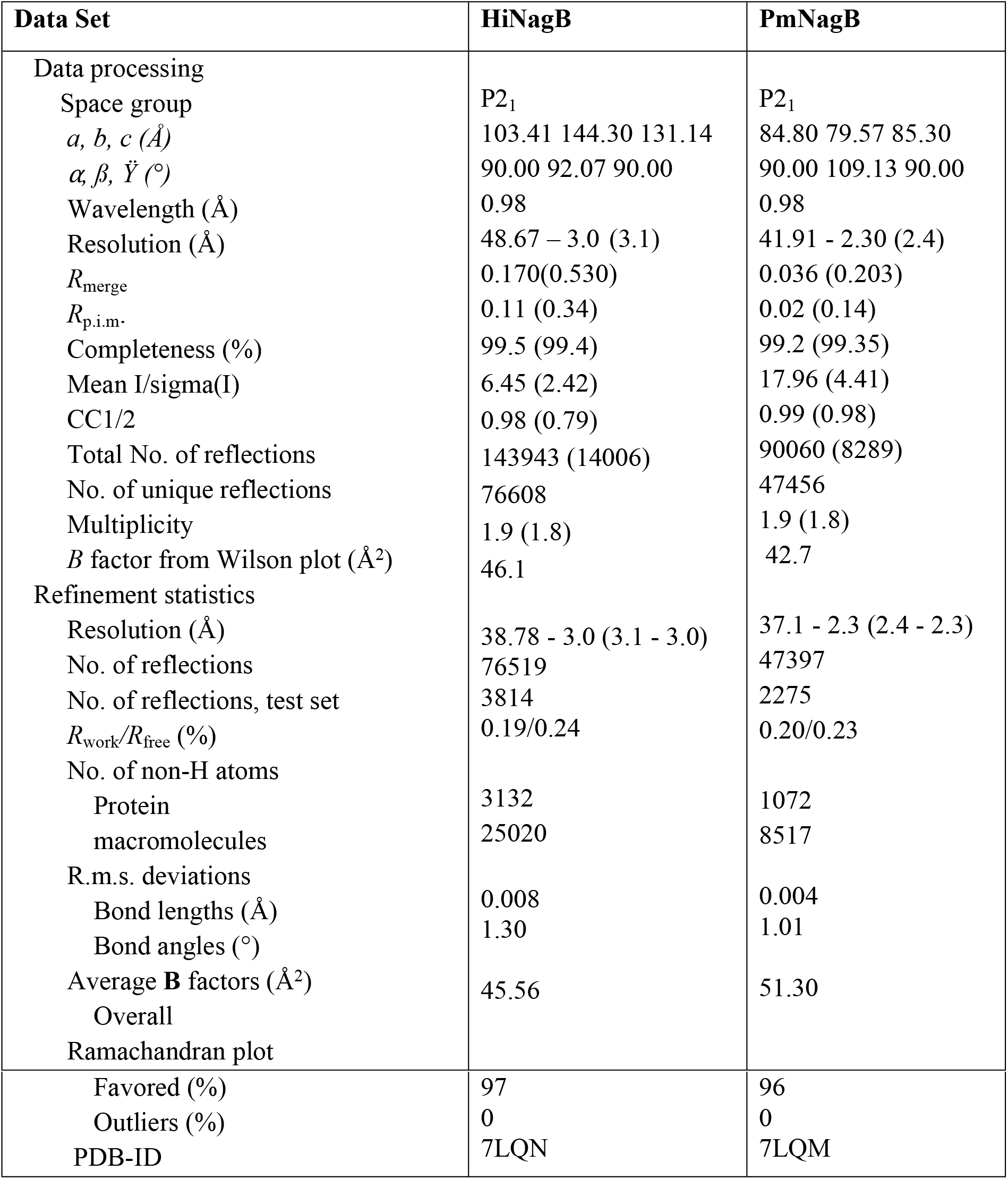
Data-processing and refinement statistics. Values in parentheses are for the highest resolution shell.

| Data Set | HiNagB | PmNagB |
| --- | --- | --- |
| Data processing |  |  |
| Space group | P2<sub>1</sub> | P2<sub>1</sub> |
| *a*, *b*, *c* (Å) | 103.41 144.30 131.14 | 84.80 79.57 85.30 |
| $\alpha$, $\beta$, $\gamma$ (°) | 90.00 92.07 90.00 | 90.00 109.13 90.00 |
| Wavelength (Å) | 0.98 | 0.98 |
| Resolution (Å) | 48.67 – 3.0 (3.1) | 41.91 - 2.30 (2.4) |
| *R*<sub>merge</sub> | 0.170(0.530) | 0.036 (0.203) |
| *R*<sub>p.i.m.</sub> | 0.11 (0.34) | 0.02 (0.14) |
| Completeness (%) | 99.5 (99.4) | 99.2 (99.35) |
| Mean I/sigma(I) | 6.45 (2.42) | 17.96 (4.41) |
| CC1/2 | 0.98 (0.79) | 0.99 (0.98) |
| Total No. of reflections | 143943 (14006) | 90060 (8289) |
| No. of unique reflections | 76608 | 47456 |
| Multiplicity | 1.9 (1.8) | 1.9 (1.8) |
| *B* factor from Wilson plot (Å<sup>2</sup>) | 46.1 | 42.7 |
| Refinement statistics |  |  |
| Resolution (Å) | 38.78 - 3.0 (3.1 - 3.0) | 37.1 - 2.3 (2.4 - 2.3) |
| No. of reflections | 76519 | 47397 |
| No. of reflections, test set | 3814 | 2275 |
| *R*<sub>work</sub>/*R*<sub>free</sub> (%) | 0.19/0.24 | 0.20/0.23 |
| No. of non-H atoms |  |  |
| Protein | 3132 | 1072 |
| macromolecules | 25020 | 8517 |
| R.m.s. deviations |  |  |
| Bond lengths (Å) | 0.008 | 0.004 |
| Bond angles (°) | 1.30 | 1.01 |
| Average **B** factors (Å<sup>2</sup>) | 45.56 | 51.30 |
| Overall |  |  |
| Ramachandran plot |  |  |
| Favored (%) | 97 | 96 |
| Outliers (%) | 0 | 0 |
| PDB-ID | 7LQN | 7LQM |

### 2.3 Structure Solution and Refinement

The phases of the *Hi*NagB and *Pm*NagB structures were obtained by molecular replacement using Phaser-MR in the PHENIX suite [15,16]. The search models were monomeric polyalanine models of PDB entry *1dea*. Model building was carried out in coot[17], and the structures were iteratively refined using the PHENIX Suite. Water molecules were automatically added during the process of refinement but were manually checked for hydrogen bonding and density fit to both (2|Fo|- |Fc|) and (|Fo| -|Fc|) maps. The structure-solution and refinement statistics are provided in Table 1.

### 2.4 Electron cryomicroscopy

Protein was purified as described for crystallography. 3 μl of *Hi*NagB at 1 mg/ml were applied to Quantifoil holey carbon grids (R 1.2/1.3, Au 300 mesh) with blotting and freezing accomplished on a Vitrobot mark IV at 18°C and 100% RH. The images of these grids showed nice isolated particles of NagB. The data of NagB were collected with Titan Krios and Falcon 3 detector in counting mode at 1.07 Å/pixel sampling with images exposed for 60 seconds with a total accumulated dose of ~30 e^-^/Å^2^ and dose fractionated into 25 frames, with each frame having a dose of ~1.2 e^-^/Å^2^. An algorithm within Relion3 was used for full-frame alignment and dose weighting[18]. The summed images were then used for automated particle picking with Gautomatch (Kai Zhang) and templates derived from manual picking, and CTF was estimated with Gift [19]. Particles were extracted with a box size of 180 pixels, and 2D classification was performed. The projections of the PDB structure were created using EMAN2 software [20]. The PDB file from the X-ray data was converted to a map file with a box size of 180 pixels and a pixel size of 1.07 Å. The map was low pass filtered to 10 Å, and the 2D projections were created for comparison.

### 2.5 MD Simulation and Analysis

We used Molecular Dynamics (MD) simulations to computationally test our hypothesis and elucidate the roles of residues in the oligomerization of NagB. The PmNagB-hexamer and *Hi*NagB-hexamer structures were subjected to molecular dynamics simulation for 50 nanoseconds using the Desmond package of Schrodinger [21,22]. Protein structures were prepared using the protein preparation wizard of Maestro (Maestro, Schrödinger, LLC, New York, NY, 2019). In System Builder, the TIP4P model was specified for water molecules, and an orthorhombic box shape was used with a buffer distance of 10 Å, followed by a minimization of the box size. The system was neutralized, and salt (NaCl) was added. The solvated system was subjected to the default relaxation protocol of Desmond before the production MD run. The relaxation protocol involves energy minimization steps using the steepest descent method with a maximum of 2000 steps. The energy minimization is done with the solute being restrained using 50 kcal/mol/Å force constant on all solute atoms and without restraints. Energy minimization is followed by short MD simulation steps which involve: 1) Simulation for 12 picoseconds at 10K in NVT ensemble using Berendsen thermostat with restrained non-hydrogen solute atoms 2) Simulation for 12 picoseconds at 10K and one atmospheric pressure in NPT ensemble using Berendsen thermostat and Berendsen barostat with restrained non-hydrogen solute atoms 3) Simulation for 24 picoseconds at 300K and one atmospheric pressure in NPT ensemble using Berendsen thermostat and Berendsen barostat with restrained non-hydrogen solute atoms 4) Simulation for 24 picoseconds at 300K and one atmospheric pressure in NPT ensemble using Berendsen thermostat and Berendsen barostat without restraints. After relaxation, production MD was run in NPT ensemble using OPLS 2003 force field [23]. For simulations, default parameters of RESPA integrator (2 femtoseconds time step for bonded and near non-bonded interactions while six femtoseconds for far non-bonded interactions) were used. The temperature and pressure were kept at 300K and 1 bar using the Nose-Hoover chain method and Martyna-Tobias-Klein method [24], respectively. The production MD was run for 50 nanoseconds. MD simulation analysis was done using the Simulation interaction diagram (SID) and Simulation event analysis (SEA) packages of the Desmond module.

## 3 RESULTS AND DISCUSSION

### 3.1 Structure of *Hi*NagB

*Hi*NagB crystallized in the P 2_1_ space group with 12 monomers in the asymmetric unit. The Matthews coefficient is 2.35 with 47.6% solvent content. The structure of *Hi*NagB was determined to 3.0 Å resolution with the final refined R-factor and R-free of 0.194 and 0.242, respectively (Table 1). The PDB ID assigned for PmNagB is 7LQN.

### 3.2 Structure of *Pm*NagB

*Pm*NagB crystallized in the P 2_1_ space group with 4 monomers in the asymmetric unit, Matthews coefficient is 2.16 with 42.6% solvent content. The structure of *Pm*NagB was determined to 2.3 Å resolution with the final refined R-factor and R-free of 0.20 and 0.23, respectively (Table 1). The PDB ID assigned for PmNagB is 7LQM.

### 3.3 Overall fold and comparison with other proteins

The monomer folds of both *Hi*NagB and *Pm*NagB are conserved and resemble the *E.coli*NagB open structure with seven-stranded parallel β sheets surrounded by eight α helices (Figure. 2A). The alpha-8 helix is also conserved in *Hi* and *Pm*, which is seen missing in the case of the monomeric deaminases from *B.subtilis* and *S. mutans* (Figure. 2B).[8,25]

**FIGURE 2.**
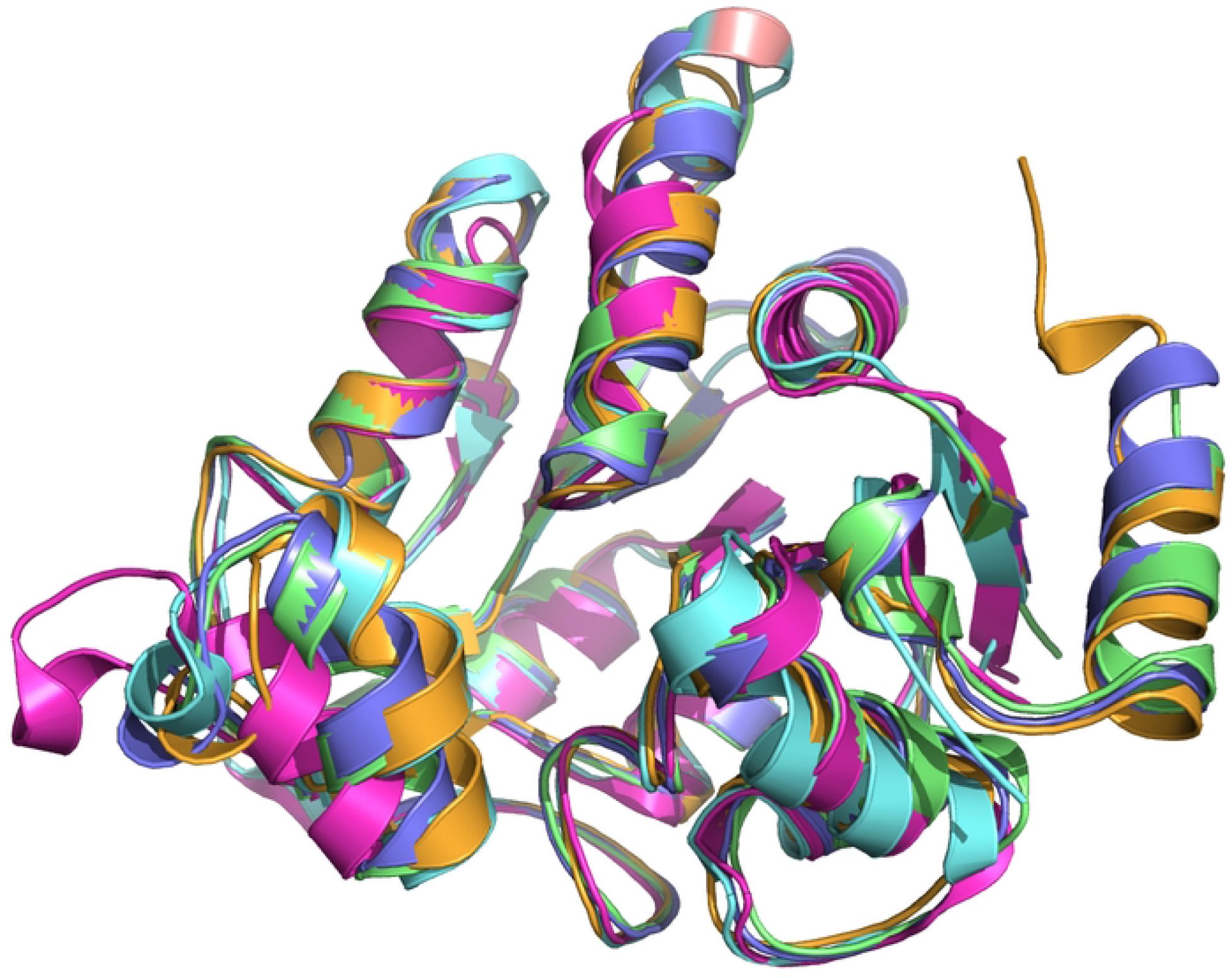
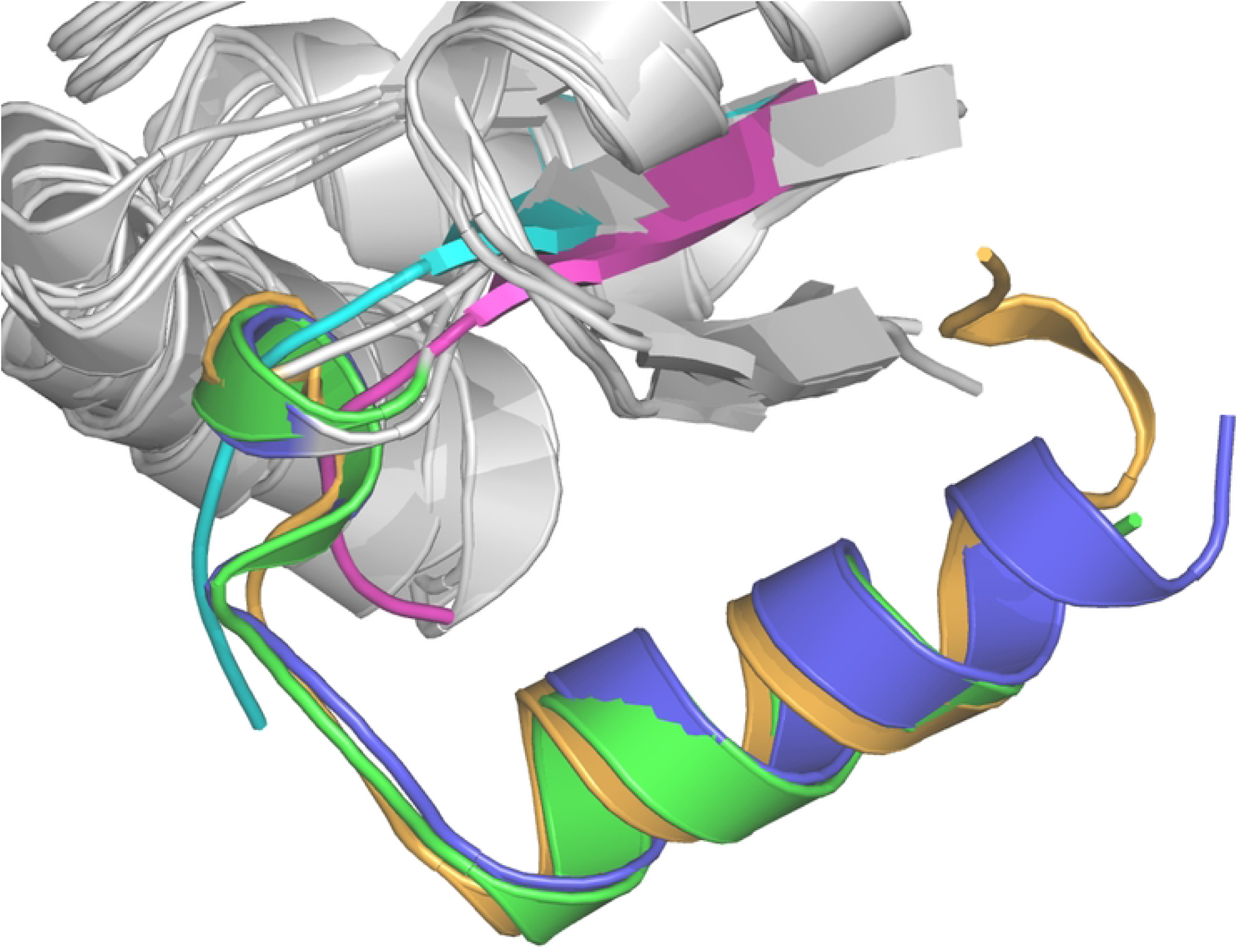
**A**: Overlay of deaminase monomers: magenta-*B.subtilis*, cyan-*S.mutans*, orange-*E.coli*, violet-*Hi and* green*: Pm*. **B**: Cartoon representation showing the presence and absence of alpha-8 helix in all the five deaminases, colour coded same as 2A.

The overall topology of the monomer resembles a modified NADH-binding domain similar to *E.coli*NagB.[11] However, in both the *Hi*NagB and *Pm*NagB, we have modified the N-terminal end with a 6-His tag. This modification hinders the formation of the allosteric binding site in the *Hi*NagB hexamer that is otherwise observed in the case of *E.coli*NagB. The allosteric site is required for GlcNAc6P to bind and allosterically regulate the enzyme.[9] However, there are no specific studies to determine the exact role of allostery on the growth rate of *E.coli*.

### 3.4 Quarternary variations - Hexameric and Dimeric deaminases

*Hi*NagB forms a hexameric structure (dimer of trimers) (Figure 3A) similar to *E.coli*NagB. The interactions between two trimers to form a hexamer are similar to that observed in *E.coli*NagB.[11] The residues in the trimeric interface are also conserved. It is important to note that even though *Pm*NagB had 80 % identity with *Hi*NagB and *E.coli NagB*, with all the active site residues and most of the interface residues conserved among the three deaminases, *Pm*NagB forms a dimer and not a hexamer (Figure. 3B).

**FIGURE 3.**
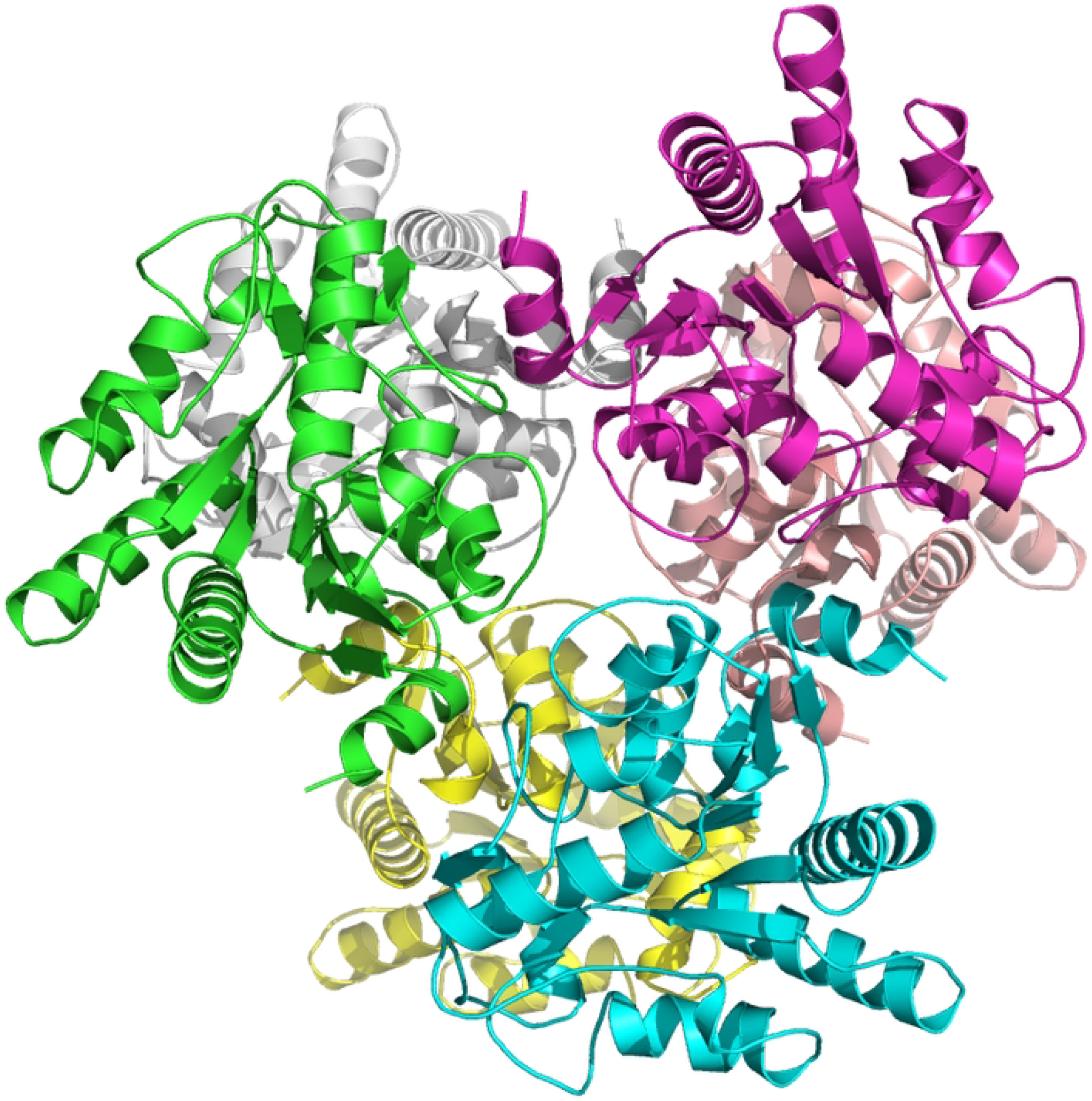
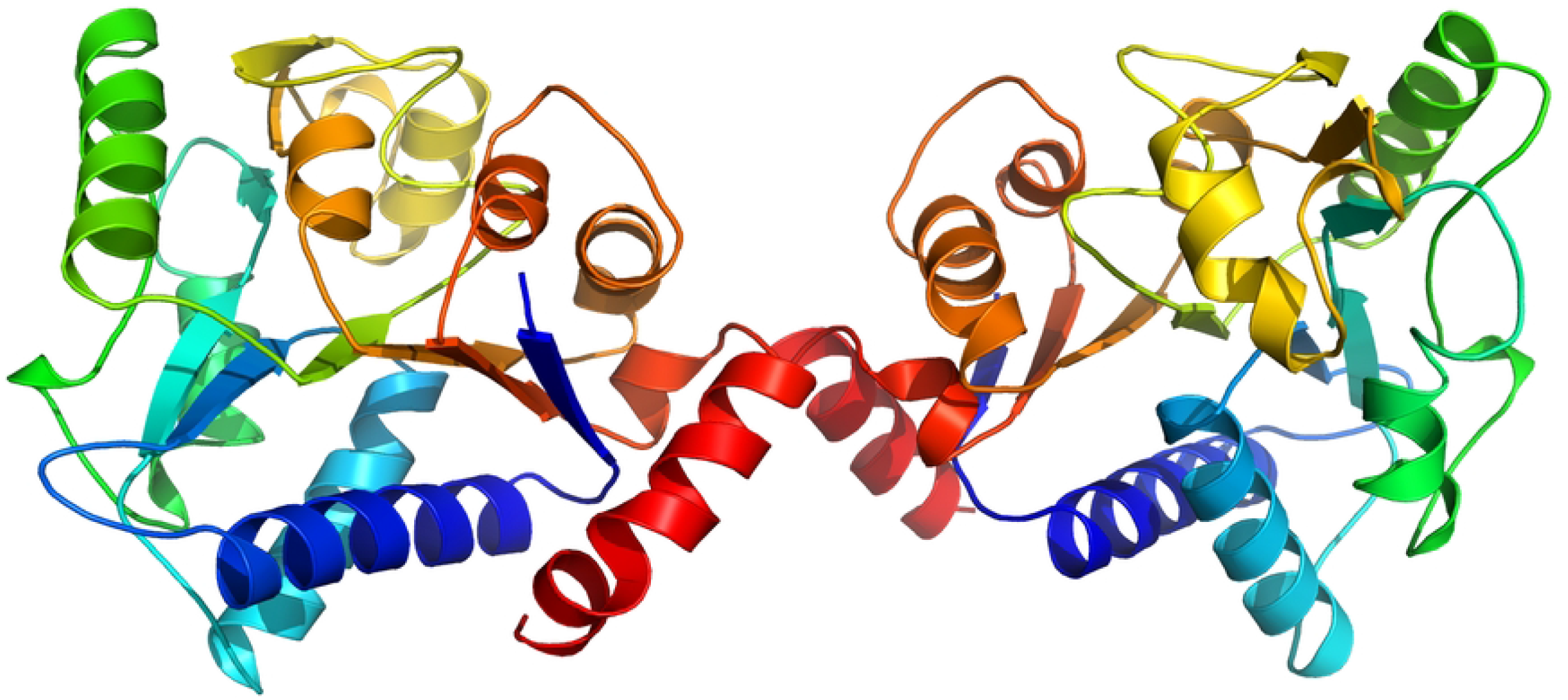
Overall structure. A: Cartoon representation of the *Hi*NagB hexamer. Each monomer is colored differently. B: Cartoon representation of the *Pm*NagB dimer. Each monomer is colored as a rainbow - with the N-terminus in Blue and the C-terminus in Red.

To ensure that the purified proteins with quaternary structures were active deaminases, we conducted steady-state kinetic studies to check the activity of the enzymes. We used ammonia Assay Kit – Modified Berthelot, Colorimetric detection from Abcam (ab102509), to study the release of ammonia. The results showed that the enzymes were active. A comparison of the K_M_ values of the deaminases revealed very similar values (Table.2). Results also showed a V_max_ of *Pm*NagB and *Hi*NagB are similar to that of *E.coli*NagB.

**TABLE 2:** Comparison of Quaternary Structure and the K_M_ of different deaminases:

| | Quaternary Structure | $K_M$ (mM) |
| --- | --- | --- |
| *E.coliNagB* | Hexamer | $0.55 \pm 0.03$[26] |
| *PmNagB* | Dimer | $0.96 \pm 0.2$ |
| *HiNagB* | Hexamer | $0.83 \pm 0.3$ |
| *B.subtilis* | Monomer | $0.13 \pm 0.02$[8] |
| *S.mutans* | Monomer | $0.21 \pm 0.03$[25] |

### 3.5 Crystal vs cryo-EM – A comparative study

We further investigated whether the hexamer seen in the *Hi*NagB crystal structure is also preserved in the solution. We made cryoEM grids and collected data, and performed 2D class average. These were then compared with the 2D projections of *Hi*NagB crystal structure with the X-ray data using the PDB file. We hypothesized that if the hexameric architecture were maintained in solution, then both the 2D class average data and the 2D projections would show near-identical features and dimensions. From an initial dataset, we manually picked 2001 particles and did 2D classification in Relion3.0.24. The raw micrograph showing isolated particles of *Hi*NagB in different orientations is shown in Figure 4A. The classes thus obtained were used as templates for automated particle picking using Gautomatch[27] 87032 particles were extracted from 565 micrographs with a box size of 180 pixels and were subjected to 2D classification. These projections correlate with the 2D averages from EM data (Figure 4B). This confirmed that the hexameric architecture of *Hi*NagB is conserved both in the crystal environment and in solution as well.

**FIGURE 4.**
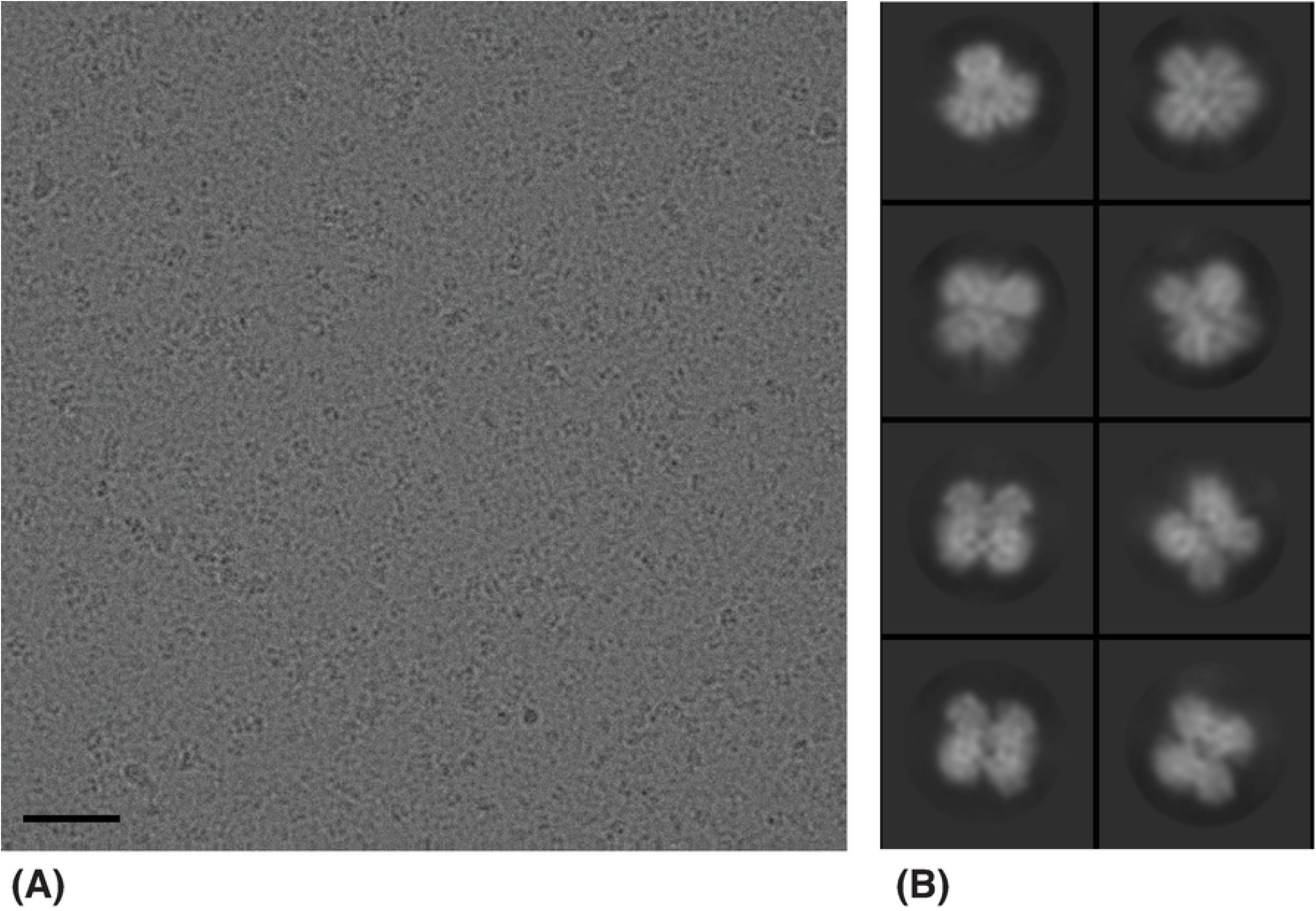
A: The raw micrograph showing isolated particles of Hi-NagB in different orientations. The particle diameter is approximately 150 Angstroms. The scale bar is 500 Angstroms. B: The image shows the 2D class averages of eight arbitrarily chosen classes. The trimeric, tetrameric, and pentameric forms can be seen

### 3.6 Bioinformatics Analysis of the interfaces

To understand the molecular basis for the observed difference in the quaternary structure of *Hi*NagB and *Pm*NagB, we investigated the dimer and the trimer interfaces of *Hi*NagB, *Pm*NagB, and *E.coli*NagB using multiple computational tools. Using one of the monomers of *Pm*NagB-dimer with structural superposition to the *Hi*NagB-hexamer, we forcefully built a hexamer model of *Pm*NagB (*Pm*NagBHexa*) to computationally understand the energetic status of such forced hexamers. Energy minimization was performed on this modeled *Pm*NagBHexa* with 100 steps of steepest descent in Swiss-PDB Viewer[28]. We calculated the interface pseudo energy for the dimer and trimer interfaces of *E.coliNagB, Hi*NagB-hexamer, and the minimized *Pm*NagBHexa* (the forcefully modeled hexamer) were calculated using PPCheck[29].

First, we analyzed the trimeric interface and observed that the interface stabilizing energy for *Hi*NagB and *E.coli*NagB was −100 kJ/mol and −170 kJ/mol, respectively. However, the calculated trimeric interface stabilizing energy for *Pm*NagBHexa* was only around −33 kJ/mol. Here, the values with less negative energies refer to poor stabilization, and values with higher negative energies refer to better stabilization, hence showing that *Pm*NagBHexa* is unstable as a hexamer. A closer look at the trimeric interface residues of *Hi*NagB and *Pm*NagBHexa*shows most of the residues were conserved, except two residues, namely Glu164 in HiNagB is Gln in PmNagB, Gln210 in HiNagB is Leu in PmNagB.

To investigate the crucial role of these two residues on trimer formation, we performed in-silico mutations of these two residues, E164Q-Q210L, in the trimeric interface of the *Pm*NagBHexa*, to those found in *Hi*NagB followed by minimization (100 sd step), and calculated the interface energy. The results suggested that the mutations did not yield any significant difference in the stabilization energies.

Simultaneously, we compared the dimer and trimer interface of *Hi*NagB and *Pm*NagB (Figure 5A and Figure 5b) to examine the contributions from the individual interface residues. The results revealed that there are around ten residues that are not conserved between the two interfaces. The stabilization energy of *Pm*NagB as a dimer is around −300 kJ/mol, whereas that of *Hi*NagB and *Pm*NagBHexa* is around −150 kJ/mol and – 80 kJ/mol. A similar trend is noticed upon analyzing the dimer interface of *Pm*NagBHexa*, *E.coli*NagB, and *PmNagB*, as shown in Figure.5B. Hence, these bioinformatic data clearly show us that *Pm*NagB is a dimer and not a hexamer similar to its *Hi* analog.

**FIGURE 5.**
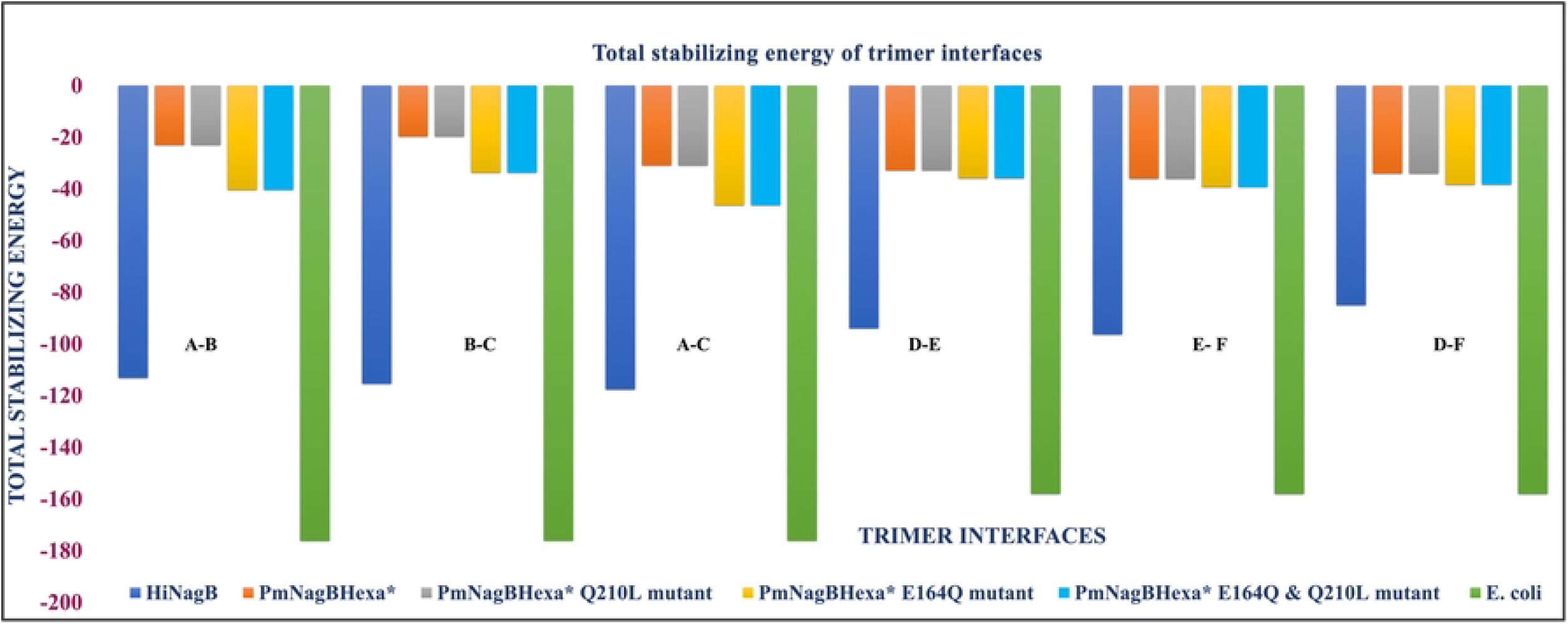
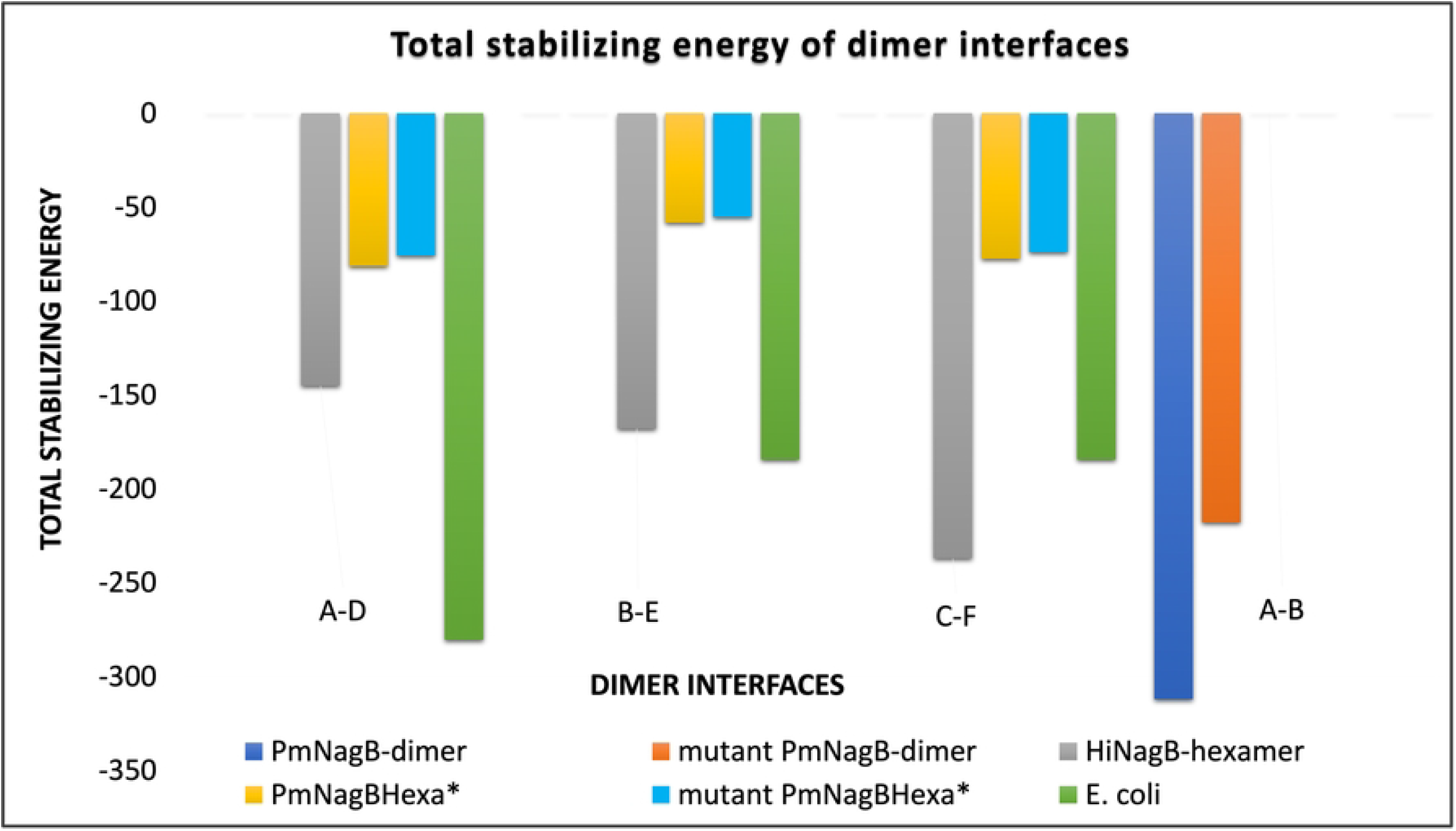
A: Total stabilizing energy of trimer interfaces of *Hi*NagB, *E.coli, PmNagB*Hexa* and its mutants (E164Q, Q210L and E164Q, Q210L double mutant) as calculated by PPCheck B: Total stabilizing energy of dimer interfaces of *Hi*NagB, *E.coli, Pm*NagBHexa* and its mutants (Y206H, P242A, T244Q triple mutant) as calculated by PPCheck.

One would normally think a molecule made of the same sequence would crystallize in a space group where the molecular symmetry coincides with the crystallographic symmetry. In the case of *Hi*NagB, the molecule is a dimer of trimers. Interestingly, the trimer is not symmetric. The angle between subunits A and B - is 120.6 °, 116.9 ° between A and C, and 122.5 ° between B and C. This results in a complete 360 ° rotation. Furthermore, the relationship is not just a pure rotation from A to B, but there is also a translation of 0.38Å along the rotation axis. In the case of A to C, the translation vector is 0.14Å, and in the case of B to C, it is 0.54Å. These are hence not just rotations but also include a translation along the axis. This should result in slight asymmetry in the interactions between the A, B, and C subunits. These are minor differences and yet are clearly visible in the attached dimplot (Figure 6). This asymmetry is not conserved among the four different trimers in the asymmetric unit. One could say that while the protein’s oligomeric state is hexameric, there is some shear between the subunits. It is currently difficult to say if this shear has an impact on the activity or the function of the enzyme. The calculations of the angles and axis were performed using the draw_rotation_axis script in Pymol. The trimer interface is shown in Figure 6.

**FIGURE 6.**
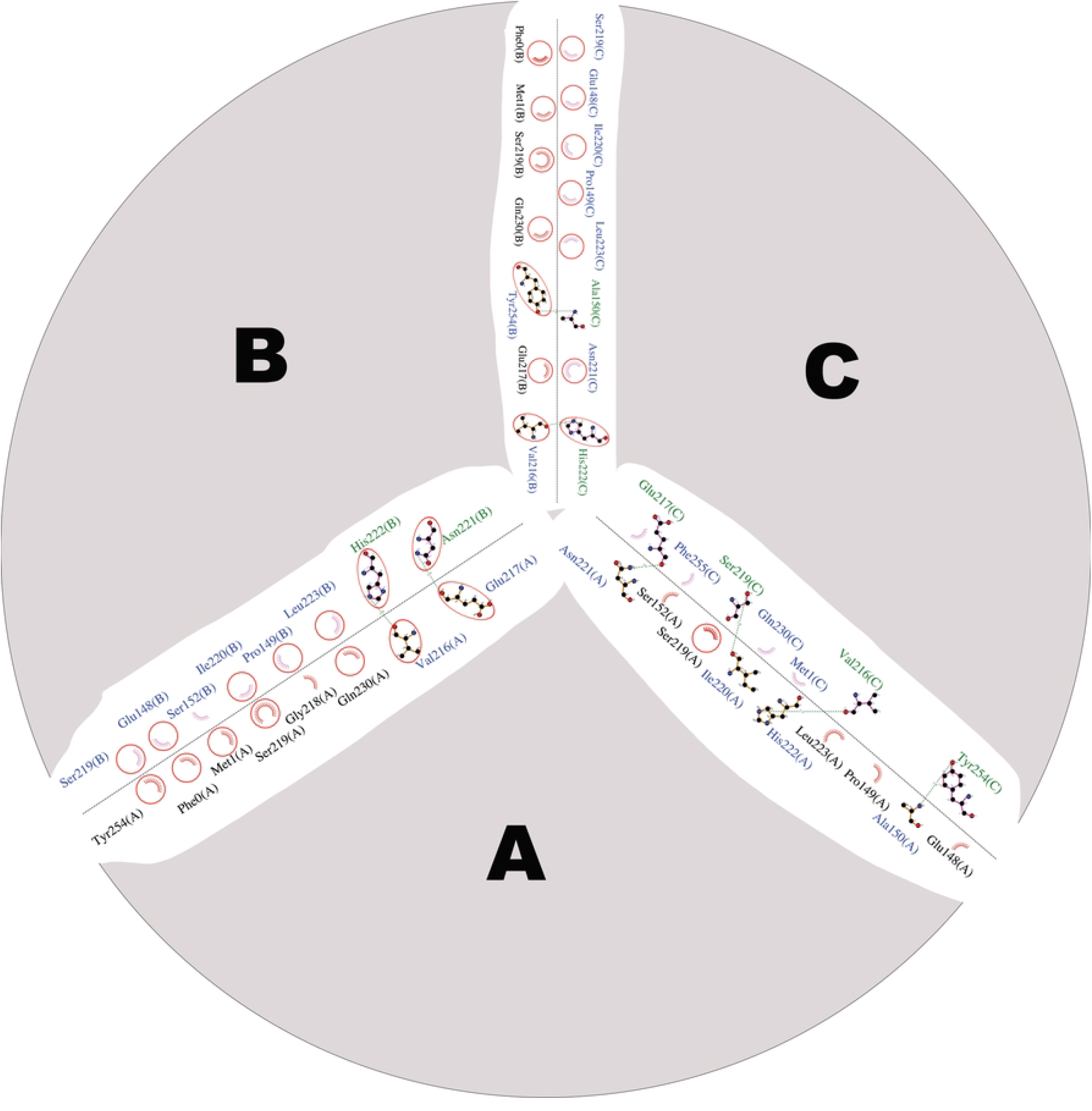
Trimer interface residue seen in *Hi*NagB using dimplot

In the case of PmNagB, the molecule is a dimer. Even though the trimer interfaces are conserved, we do not observe the interaction seen in the case of *Hi*NagB and *E.coli*NagB.

Figure 7 shows the interface interactions that favor dimerization. This interface plays a crucial role in the quaternary state of the protein. The dimer interface residues might serve as a signature motif between the monomeric, dimeric, and hexameric family of deaminases that can help in predicting the quaternary structure of other deaminases. Quaternary variations are often either related to allostery or stability. The presence of allostery in some of the enzymes would suggest that is probably the role. An important question that is yet to be answered is the role of these quaternary variations or allostery in the physiological function of these proteins.

**Figure 7.**
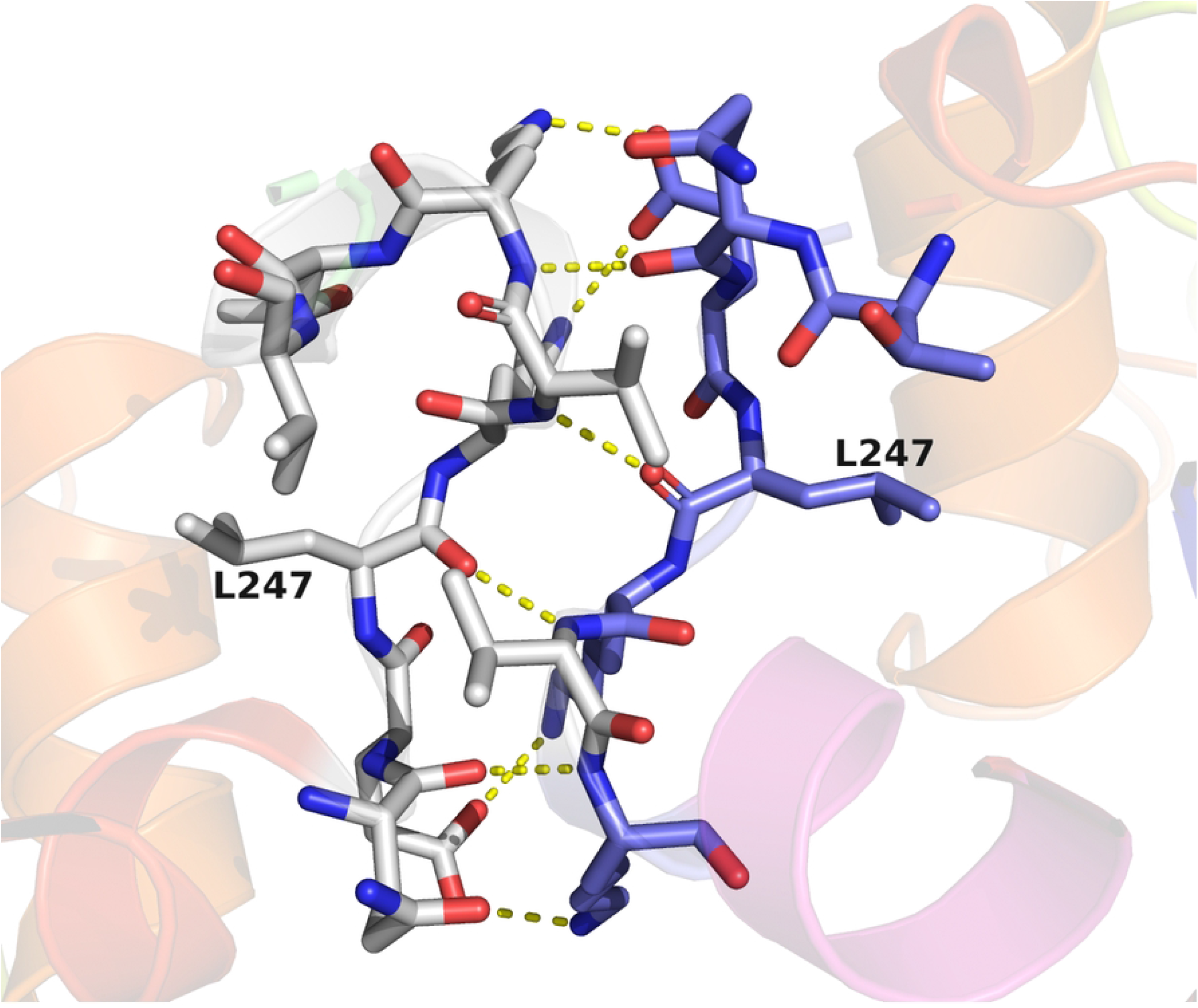
The conserved dimeric interface. Here the interface is drawn from the structure of PmNagB.

## ACKNOWLEDGMENTS

SS would like to thank DBT/Wellcome Trust India Alliance for the generous funding through the Early Career fellowship. We thank SOLEIL Synchrotron (Proxima-1 beamline) for providing the beamtime for data collection. We would like to thank the NCBS cryoEM Facility. We would like to thank Dr. Sucharita Bose for her help with the initial data processing and guidance during the writing of the manuscript.

## Data Availability

The two protein structures reported here have been deposited in the protein data bank with IDs of 7LQN and 7LQM.

The authors declare that they do not have any competing interests.

